# T-RHEX-RNAseq – A tagmentation-based, rRNA blocked, random hexamer primed RNAseq method for generating stranded RNAseq libraries directly from very low numbers of lysed cells

**DOI:** 10.1101/2022.10.20.513000

**Authors:** Charlotte Gustafsson, Julia Hauenstein, Nicolai Frengen, Aleksandra Krstic, Sidinh Luc, Robert Månsson

## Abstract

**Background:** RNA sequencing has become the mainstay for studies of gene expression. Still, analysis of rare cells with random hexamer priming – to allow analysis of a broader range of transcripts – remains challenging.

**Results:** We here describe a tagmentation-based, rRNA blocked, random hexamer primed RNAseq approach (T-RHEX-RNAseq) for generating stranded RNAseq libraries from very low numbers of FACS sorted cells without RNA purification steps.

**Conclusion:** T-RHEX-RNAseq provides an easy-to-use, time efficient and automation compatible method for generating stranded RNAseq libraries from rare cells.

## BACKGROUND

The field of transcriptomics has during the last two decades evolved from analysis of select genes with PCR-based methods, to broad analysis using microarrays and finally, to the now standard use of RNA sequencing (RNAseq) to characterize full transcriptomes [1]. Principally, RNAseq is achieved by reverse transcribing RNA into cDNA, introducing sequencing adapters and applying next-generation sequencing to determine the presence of transcripts.

Through this process, the strand-of-origin information can be maintained using different strategies, allowing for accurately resolving sense and antisense transcription [2, 3]. A common strategy is the incorporation of deoxy-UTP (dUTP) during second strand cDNA synthesis [4], followed by a digestion of the uracil containing strand before amplification of the library.

The sequencing of full length polyadenylated RNA – using oligo-dT priming – has been the mainstay for RNAseq. However, better understanding of RNA biology has come with increased interest in studying also other RNA species or incompletely processed RNA lacking polyadenylation [1]. The use of random hexamer priming allows for analyzing a broader range of RNA species, but this comes with the drawback that the ribosomal RNA (rRNA) – that can constitute up to 90% of total RNA – is reverse transcribed together with the RNA of interest. This makes a reduction of rRNA representation in the library an attractive strategy for limiting the need for extensive sequencing. Several strategies to restrict inclusion of rRNA have been developed, including depletion of rRNA sequences [5–7], enzymatic degradation of rRNA [8, 9] or simply blocking of rRNA from undergoing reverse transcription [10].

Early RNAseq libraries were generated by ligation of sequencing adapters to double stranded (ds) cDNA in a multi-step process. The use of transposase (Tn5) to fragment ds DNA and integrate sequencing adapters in a single step – so called tagmentation – was first described by Adey et al., [11]. Because of the simplicity, tagmentation has seen wide use in genomics applications ranging from whole genome sequencing [12] to ATACseq [13] and ChIPseq [14, 15]. In collaboration with Epicenter, Gertz et al., [16] published the Directional Tn-RNAseq method and demonstrated that tagmentation in combination with the use of dUTP incorporation could be used to generate stranded RNAseq libraries. In a refined version, this method was briefly made commercially available from Epicenter as the TotalScript kit. The kit provided a straightforward procedure for generating random hexamer primed stranded RNAseq libraries from 1-5ng of total RNA, although an effective solution for reducing the rRNA contribution to the library was lacking. Analogous to how stranded RNAseq libraries were generated using forked adapters and dUTP incorporation [4], the Directional Tn-RNAseq method relied on introducing the i5 adapter on the 5’ side of the transposed cDNA. This was achieved by introducing only the i5 adapter in the tagmentation step. The i7 adapter was subsequently introduced by replacing the part of the i5 adapter that is not covalently linked to the tagmented DNA with an i7 adapter oligo. Noteworthily, this allows for sequencing all tagmented molecules, in contrast to standard tagmentation where only half the library fragments acquire the i5-i7 combination needed for sequencing. With the PCR amplification of the library being performed after the tagmentation step, the protocol further allowed for using the recurrence of the same Tn5 integration sites to estimate and compare the potential contribution of biased PCR amplification to the library.

The current market offers several commercially available kits for generating RNAseq libraries from ≤1ng of RNA, including the SMARTer Stranded total RNA-seq Pico kits (Takara), NEBNext Single cell/Low input RNA prep kit (New England Biolabs; NEB), and Ovation SoLo RNAseq preparation kit (Tecan). However, none of these offers a solution with the combinatorial use of cell lysates as input for the cDNA synthesis, random hexamer priming, and effective removal of rRNA, while at the same time taking advantage of the streamlined workflows that can be achieved using tagmentation for RNAseq [17–20].

To establish a protocol that could take advantage of this combination to analyze a wide spectrum of RNA in very low numbers of FACS sorted cells, we revisited the use of tagmentation with one adapter to achieve stranded RNAseq libraries [16]. Based on the re-creation and refinement of this approach, we here present a tagmentation-based, rRNA blocked, random hexamer primed RNAseq method (T-RHEX-RNAseq) for generating stranded RNAseq libraries from cells without prior RNA purification. T-RHEX-RNAseq is technically straightforward, time efficient, compatible with automation and can be performed on input material ranging from nanograms of purified RNA down to very low numbers of FACS sorted cells.

## RESULTS

### Re-establishing directional tagmentation-based RNAseq library construction

The Directional Tn-RNAseq protocol by Gertz et al., [16] in brief relied on synthesis of ds cDNA with dUTP incorporation in the second strand, tagmentation with i5 compatible adapters alone, introduction of the i7 adapter by oligo replacement, digestion of the dUTP containing strand and amplification of the library followed by sequencing.

Through a series of pilot experiments on purified total RNA, we devised a similar strategy for generating stranded RNAseq libraries using commercially available reagents (data not shown). Together, this formed the core steps of the protocol outlined in Figure 1. In brief, the NEBNext Ultra II RNA First Strand Synthesis module (NEB) in combination with the NEBNext Ultra II Directional RNA Second Strand Synthesis module (NEB) was used to generate ds cDNA (Fig. 1iii-iv). These reagents were chosen because of the usage of random hexamer priming, stated low input need (1-100ng prepared RNA) and incorporation of dUTPs in the second strand synthesis. Following second strand synthesis, Tn5 with only i5 compatible adapters was used to tagment the ds cDNA (Fig. 1v and S1). Subsequently, oligo replacement was used to introduce the i7 adapter (Fig. 1vi-vii and S1). For this process to work, the annealed (index containing) i7 replacement oligo needs to be covalently attached to the free 3’end left by the tagmentation. To achieve this, we used Sulfolobus DNA Polymerase IV (NEB) – which lacks both strand displacement and 5’-3’ exonuclease activity which otherwise could displace or degrade the annealed i7 oligo – in combination with E. coli DNA Ligase (NEB). Together, these enzymes fill the gap left by the tagmentation and seal the remaining nick between the 3’end of the tagmented ds cDNA and the i7 oligo (Fig. 1vii and S1). Finally, to generate sequencing compatible libraries, we used the Phusion polymerase together with custom i5 completion primers and i7 amplification primers (Fig. 1viii and S1).

**Figure 1.**
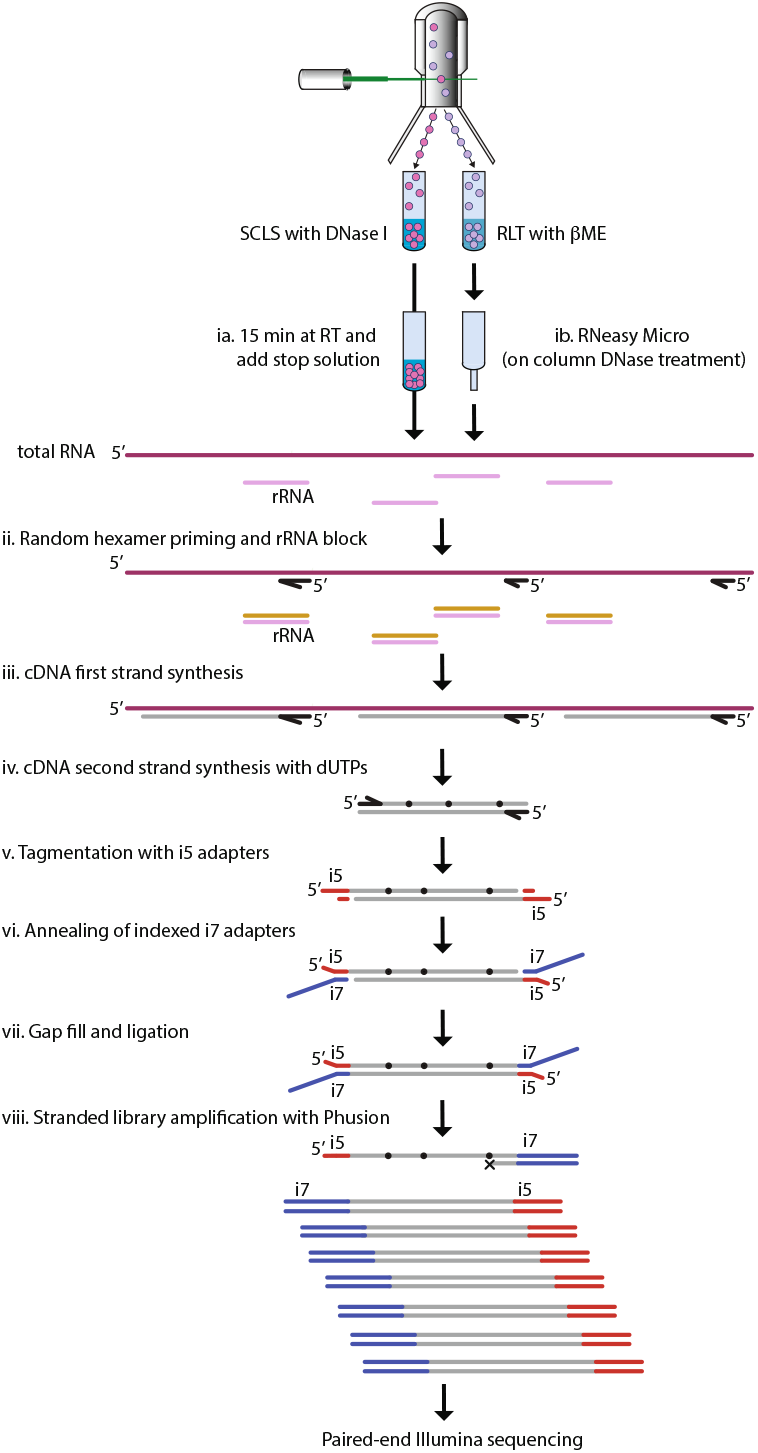
Schematic outline of the T-RHEX-RNAseq protocol. In brief, lysed cells (ia) or total RNA (ib) are subjected to random hexamer priming with rRNA blocking (ii), reverse transcription (iii) and second strand synthesis with dUTP (iv). Subsequently, tagmentation (v) and oligo replacement (vi-vii) are used to introduce adapter sequences before the library is amplified (using dUTP incompatible Phusion) to generate stranded libraries (viii). For details on adapter sequences and primers see Figure S1.

### Stranded RNAseq libraries can be generated from very low numbers of FACS sorted cells when RNA purification and dUTP-based degradation steps are omitted

To refine the protocol from the pilot experiments and make it feasible to use low numbers of FACS sorted cells, we next tested if we could omit the RNA purification step and instead directly prepare libraries on lysed cells. At the same time, we wanted to validate that we could simplify the protocol and generate stranded libraries without enzymatically degrading the dUTP containing strand. The rationale behind this being that the High-Fidelity Phusion polymerase (NEB) should generate stranded libraries during the amplification process simply through the inability of the enzyme to read dUTP in the template [21].

To this end, we FACS sorted a series of progressively lower numbers of cells (500, 250, 100 and 50 MM1.S cells) directly into Single cell lysis solution (SCLS, Invitrogen) with DNase I – to facilitate cell lysis and degradation of genomic DNA – and prepared libraries on this material as outlined in Figure 1. Inspection of the mapped reads showed that expression originating from polyadenylated (*Irf4*, *Gnl3*, *Glt8d1* and *Xist*) and non-polyadenylated (*Neat*) [22] genes as well as intronic regions of incompletely spliced transcripts (as seen for *Irf4*) could readily be observed, as expected from the use of random hexamer priming (Fig. 2A). The clear localization of reads to gene bodies indicated that no overt DNA contamination was found in the libraries and hence that the DNase treatment, done as part of the cell lysis, was adequate to hinder genomic DNA from contributing to the library. In addition, the genome browser tracks suggested that transcriptional profiles were maintained even in the samples generated from the lowest cell numbers. Confirming this notion, library quality was maintained across the full range of input cells with all samples globally having highly similar transcriptome profiles (correlation >0.98) (Fig. 2B). The libraries further maintained comparable duplication rates (Fig. 2C), mappability (Table S1) and library insert sizes (Fig. 2D).

**Figure 2.**
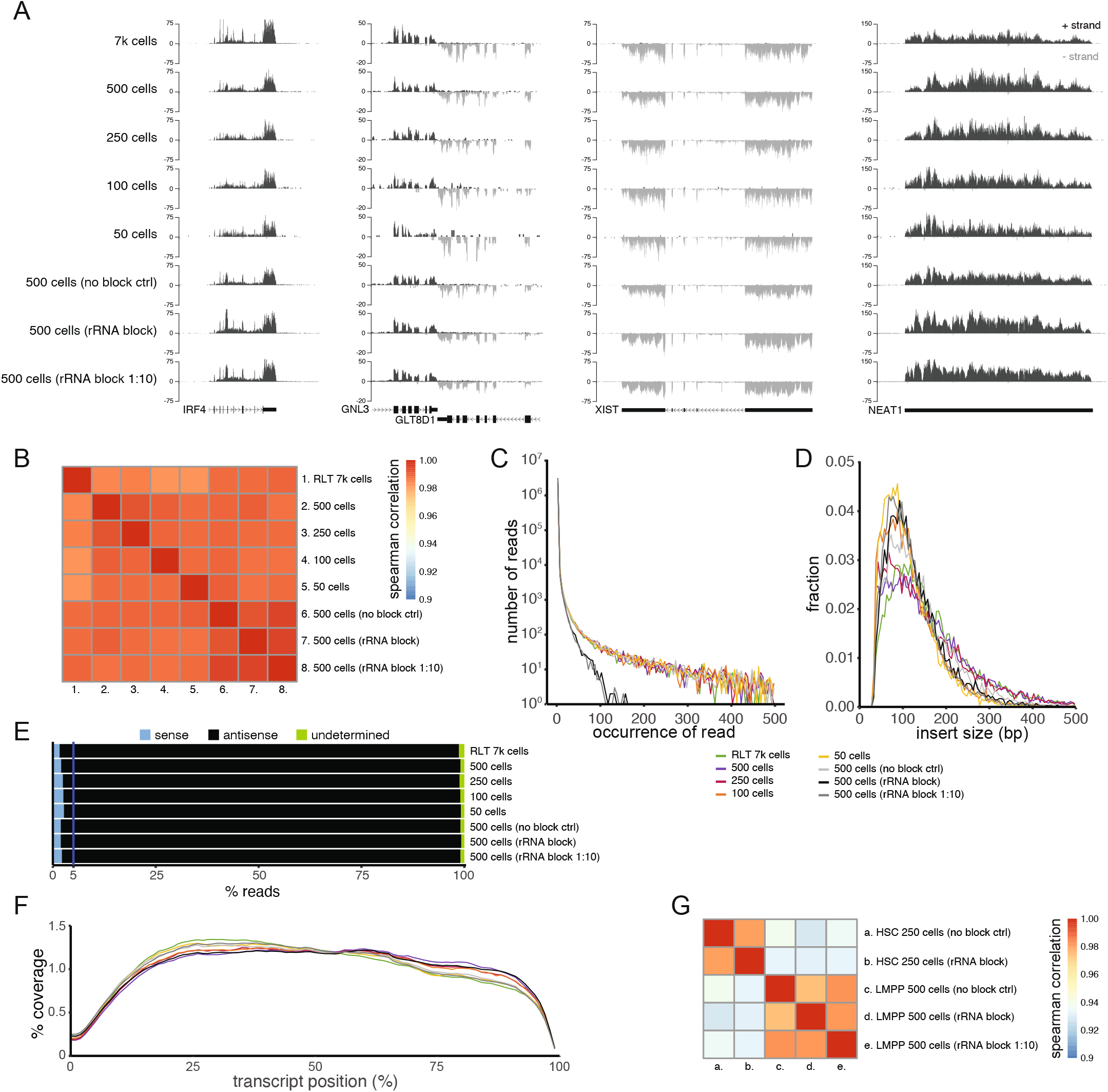
Pilot experiments and establishment of the T-RHEX-RNAseq protocol. (A) Tracks showing reads on the plus and minus strand in the *Irf4*, *Gnl3*, *Glt8d1*, *Xist* and *Neat1* genomic regions. RNAseq libraries were prepared using purified total RNA (7k MM1.S cells) or directly from the indicated numbers of MM1.S cells lysed in Single cell lysis solution (SCLS). The use of rRNA blocking reagents and dilution of the blocking reagent is indicated in parenthesis. (B) Spearman correlation between rlog of gene expression in MM1.S samples. (C) Reoccurrence (duplication rates) of reads in the indicated libraries. (D) Library insert size distributions. (E) Percentage of reads being antisense (stranded), sense (unstranded) or undetermined (in regions with overlapping antiparallel transcripts) in exons. (F) Mean distribution of coverage (%) across the length of all transcripts. (G) Spearman correlation between rlog of gene expression in primary mouse hematopoietic stem cells (HSCs) and lymphoid primed multipotent progenitors (LMPPs).

In line with the libraries being stranded – even without dedicated steps to degrade the dUTP containing strand – tracks clearly showed directional transcription (Fig. 2A). A very high enrichment of stranded reads was observed for genes without a close antiparallel promoter such as *Irf4*, *Xist* and *Neat1*, that respectively showed 182-, 215- and 205-fold enrichment of antisense reads (>99.4% antisense reads on average) over sense strand reads. Genome-wide, we found that <2.7% of reads in exons were called as non-stranded (sense) reads (Fig. 2E).

Random hexamer priming should provide a relatively even coverage across the gene body. Analyzing this, we found that overall, the coverage was substantially even, though depletion was observable in the first 15% of the 5’-end and in the outermost 3’-end (Fig. 2F). This loss of coverage in the 5’-end can be attributed to the need for Tn5 to integrate in the very outermost part of the 5’-end for it to be represented in the library. Presumably, this effect is mitigated in the 3’-end by the Tn5 integrating in the polyA-tail outside of the transcript specific sequences.

Overall, we concluded that stranded RNAseq libraries could be generated directly from the lysate of very low numbers of FACS sorted cells.

### rRNA can effectively be eliminated from the library by blocking prior to reverse transcription

While our initial libraries gave a good representation of expressed RNA, the rRNA contribution to the libraries was substantial in our test experiments (on average 63% of reads) (Table S1). With the rationale that reducing the rRNA contribution to the library before reverse transcription would be preferable to lower the need for deeper sequencing and improve representation of non rRNA molecules in the library, we tested the QIAseq FastSelect -rRNA HMR kit. While proprietary, the reagent presumably constitutes oligos that block the rRNA molecules from undergoing reverse transcription. To incorporate this in our protocol, we added the QIAseq FastSelect -rRNA HMR reagent at the random hexamer annealing step (Fig. 1ii) and replaced the conventional annealing with stepwise decreases in temperature from 75°C to 25°C as per the QIAseq FastSelect protocol. We found that this reduced the rRNA contribution from an average of 63% (range 60-86%) to <1% of reads (Table S1). Compared to unblocked samples, libraries otherwise maintained overall profiles of mapped reads (Fig. 2A) as well as similarity on the whole-transcriptome level (correlation >0.98) (Fig. 2B). The rate of duplication also markedly decreased in the blocked samples (Fig. 2C), indicating that the elimination of the rRNA improved the representation of original non-rRNA molecules in the library.

With the inclusion of the rRNA blocking at the reverse transcription stage, this formed the basis for our stable version of the T-RHEX-RNAseq protocol (see Materials and Methods and supplemental working protocol).

### T-RHEX-RNAseq can be performed on rare primary cells with high reproducibility

As the T-RHEX-RNAseq protocol was established using the MM1.S cell-line, we next wanted to validate that the protocol could be used to generate libraries on small, mainly resting primary cells that can be expected to contain significantly less RNA per cell. We did this by generating libraries from 250 or 500 FACS sorted primary hematopoietic stem cells (HSCs) and lymphoid primed multipotent progenitors (LMPPs) from mouse bone marrow. Visual inspection of mapped reads indicated that high quality stranded libraries were generated (Fig. S2A) and high correlation was found between replicates (correlation >0.97) (Fig. 2G). These experiments further supported the conclusion that the overall transcriptional profiles remained unaffected by the inclusion of the rRNA blocking reagent (Fig. 2G) and that blocking the rRNA resulted in improved representation of other RNA species to the library as indicated by the lower duplication rates (Fig. S2B). Further analysis showed that the primary cell libraries displayed high mapping efficiency (>74%) (Table S1) and low rRNA contribution (average ≤2.2%) in blocked samples (Table S1).

The experiments performed to get proof of principle and establish the T-RHEX-RNAseq method showed that the method could generate data from rare cells. However, the number of samples analyzed was relatively small. To address the reproducibility, we reanalyzed the T-RHEX-RNAseq proof of principle data from the primary cells together with our subsequently generated T-RHEX-RNAseq data from antigen specific CD4 T cells (1000 cells) [23] and hematopoietic stem/progenitor cells (HSPCs) (250-500 cells) [24]. This showed that reproducibility was very high both within the published data sets (correlation >0.98 and >0.95 for the T cells and HSPCs respectively) as well as between the HSPC samples from the proof-of-principle experiments and corresponding cells in the subsequently generated hematopoietic stem/progenitor data (correlation >0.97) (Fig. S3).

Overall, we conclude that directional T-RHEX-RNAseq allows for reproducible generation of high-quality RNA expression data from very low numbers of FACS sorted primary cells.

## DISCUSSION

We have here described stranded T-RHEX-RNAseq, an effective, streamlined and highly reproducible workflow for preparing stranded RNAseq libraries from very low numbers of cells. The simplicity of the protocol, at the core, rests on the use of transposase (loaded with only i5 adapters) in combination with dUTP incorporation to generate the library and maintain strand specificity [16]. With a process analogous to that described by Gertz et al., we could effectively prepare libraries from ng amounts of total RNA. However, the direct use of this protocol had limitations for broader implementation including the substantial rRNA contribution to the library caused by the random hexamer priming and losses of material associated with bead cleanup steps and, in particular, RNA purification.

As the losses associated with RNA purification set direct limits on the lowest possible input cell number, the key to being able to perform RNAseq on low cell numbers – rather than on low amounts of RNA purified from often substantially higher amounts of cells – is removing the need for RNA purification. Aiming to establish the T-RHEX-RNAseq protocol, we effectively removed this limitation by directly preparing the libraries on lysed cells. This allowed us to generate libraries with maintained quality from as little as 50 cells. Considering the maintained quality with these low input cell numbers, it is feasible that even fewer or also single cells could be analyzed using this protocol. The latter in particular, if barcodes could be introduced at the tagmentation step [25], which would allow for pooling samples prior to the first bead-cleanup step. While we did not specifically investigate the impact of removing the dUTP degradation and the associated bead cleanup step (prior to library amplification), this likely improved the ability of the T-RHEX-RNAseq protocol to generate libraries on low cell-numbers while maintaining more than adequate strandedness.

While the use of random hexamer priming on total RNA allows for analyzing a wider range of transcripts, it comes with the disadvantage that rRNA is effectively reverse transcribed. The significant contribution of rRNA to the library seen in our initial libraries hence was a significant hindrance from sequencing RNA of interest in a cost-effective manner. Given the scarce input material and the complexity of depleting the rRNA derived part of the final library, we reasoned that blocking rRNA from undergoing reverse transcription would by far be the simplest and most effective solution to implement. In the T-RHEX-RNAseq protocol, we found that this approach substantially mitigated the issue of rRNA contributing to the library. This not only lowered the sequencing needed to acquire adequate depth for analyzing RNA expression but interestingly also improved the representation of RNA of interest in the library. The latter suggests that this approach of blocking rRNA before reverse transcription could be generally beneficial for RNAseq approaches using cell lysates or total RNA as starting material.

In terms of implementation, the T-RHEX-RNAseq offers flexible use of purified RNA and cells collected in SCLS. Practically, storing cells in SCLS in individual PCR tubes – that can be attached to a holder to generate a 96-well plate – offers an adaptable solution where samples can easily be selected for analysis and collected into larger batches that can then be processed in semi high-throughput manner in 96-well format. While we have commonly generated T-RHEX-RNAseq libraries over two days, it is practically possible for an experienced user to generate libraries in a single day and then perform sequencing on the NextSeq 500/550/2000 systems the following day. This makes it possible to start processing the sequencing data less than 48h after the start of the library preparation which overall makes this a time efficient process. In terms of performing cost efficient sequencing, the smallest versions of NextSeq sequencing kits (75 or 50 cycles depending on system) are both sufficient for paired-end 41 sequencing, which gives good coverage for most library inserts. While longer reads likely would improve mapping efficiency, this provides a good trade-off between data quality, cost and time with the sequencing taking approximately 12h.

Hence, overall T-RHEX-RNAseq offers an easily implemented and convenient solution for generating stranded RNAseq libraries from total RNA down to very low numbers of FACS sorted cells.

## CONCLUSION

We here describe T-RHEX-RNAseq, an easy-to-use protocol for generating stranded RNAseq libraries from low numbers of FACS sorted cells. The protocol uses the simplicity offered by tagmentation and takes advantage of random hexamer priming while avoiding limitations caused by rRNA removal and RNA purification steps. This offers a time efficient and reproducible method for analyzing complete transcriptomes, that is compatible with applications ranging from analysis of rare cells to large scale projects requiring automation.

## METHODS

### Cells and FACS sorting

MM1.S was cultured in RPMI-1640 with GlutaMAX and HEPES supplemented with 10% fetal calf serum and 1% penicillin-streptomycin. Defined numbers of MM1.S cells were FACS sorted on a FACSARIA IIu (BD Biosciences) using propidium iodine to exclude dead cells. Primary mouse HSCs (LIN^−^ SCA1^+^ KIT^+^ CD48^−^ CD150^high^) and LMPPs (LIN^−^ SCA1^+^ KIT^+^ FLT3^high^ CD150^−^) were FACS sorted from bone marrow as previously described [24]. To perform RNAseq on lysed cells, cells were FACS sorted directly into 5 or 10μl Single cell lysis solution (SCLS; Single cell lysis kit Invitrogen, cat# 4458235) with DNase I according to manufacturer’s instructions with the exception that the lysis reaction was incubated for 15min at room temperature (RT) before adding stop solution. To perform RNAseq on purified total RNA, cells were FACS sorted into RLT buffer with β-mercaptoethanol. RNA was extracted using the RNeasy Micro Kit (Qiagen, cat#74004) with on-column DNase I treatment according to manufacturer’s instructions. Both samples in SCLS and RLT were snap frozen on dry ice and stored at −80°C until use.

### Tn5 with i5 adapters

Tn5 loaded with only i5 compatible adapters was kindly provided by Prof. Rickard Sandberg or generated by loading purified un-loaded Tn5 tagmentase (Diagenode cat#C01070010) with adapters according to the manufacturer’s instructions (see Fig. S1 and working protocol for adapter sequences).

### T-RHEX-RNAseq

Double stranded cDNA was prepared from lysed cells in SCLS or purified total RNA using NEBNext Ultra II RNA First Strand Synthesis (NEB cat#E7771) and NEBNext Ultra II Directional RNA Second Strand Synthesis (NEB cat#E7550) modules according to manufacturer’s instructions. In rRNA blocked samples, QIAseq FastSelect -rRNA HMR kit reagent (Qiagen cat# 334386) was added to the first strand cDNA synthesis reaction according to manufacturer’s instructions with the exception that only one tenth the amount was used for the majority of libraries. Stranded library preparation was subsequently initiated by adding tagmentation buffer (50mM tris acetate, 25mM Mg acetate, 50% dimethylformamide) and Tn5 with i5 adapters to ds cDNA. After 5 min at 55°C, the reaction was stopped by incubating with 5μl 1% SDS for 5min at room temperature and samples were purified using AmPure XP beads at a ratio sample:beads of 1:1.2 and eluted in water. Introduction of indexes through oligo replacement [16] was carried out by adding custom made i7 replacement oligos and gap fill buffer (165mM tris acetate, 330mM potassium acetate, 50mM Mg acetate, 2.5mM DTT, 1mM beta-NAD and 1.25mM each of dATP, dCTP, dGTP and dTTP). After 30min incubation at 37°C, gap fill enzymes were added (1U Sulfolobus DNA Polymerase IV NEB cat# M0327S and 10U E. coli DNA Ligase NEB cat# M0205L) and incubation continued for an additional 30min at 37°C. Samples were purified using AmPure XP beads at a ratio sample:beads of 1:1.2 and eluted in 12μl water. Libraries were subsequently PCR amplified using Phusion HF PCR Master Mix (NEB) with the following PCR program: 95°C 2 min; followed by 16 cycles of 94°C 10 s, 60°C 30 s and 72°C 1 min. Library cleanup was done using AmPure XP beads at a ratio sample:beads of 1:0.9. DNA concentrations in purified samples were measured using the Qubit dsDNA HS Kit (Invitrogen). Barcoded libraries were pooled and paired-end sequenced (2×41 cycles) using the Illumina platform (NextSeq500, Illumina).

### Analysis of RNAseq data

Samples were analyzed using the nf-core/rnaseq pipeline (v.3.3) [26, 27] with the --remove_ribo_rna option and the default rRNA database. Mouse and human samples were aligned to the mm10 and hg38 reference genome, respectively. Mapping rates and rRNA contribution to the libraries were derived from nf-core via MultiQC [28]. For visualization of read coverage, tracks obtained from nf-core were normalized to 10^7^ mapped reads and uploaded to the UCSC genome browser [29]. To perform correlation analysis between samples, per gene read counts were derived from nf-core, read counts of expressed genes (≥30 reads in ≥3 human or mouse samples) normalized using rlog transformation, and the spearman correlation was calculated. To compare duplication rates between samples, bam files were downsampled to the same number of mapped reads and duplication rates determined using the read_duplication.py function from the RSeqQC package (v.2.6.4). For visualization, the occurrence of duplicated reads was calculated for each 5-step increment and plotted using R (v4.1.1) and the ggplot2 package (v3.3.6). Library insert sizes were derived from nf-core, binned into 5bp bins and plotted using ggplot2. Inferred strandedness was determined using the RSeqQC package considering reads overlapping known exons. Coverage across the length of the gene body was derived from nf-core, re-calculated to % of total coverage and plotted using ggplot2.

## DECLARATIONS

### Ethics approval and consent to participate

Not applicable.

### Consent for publication

Not applicable.

### Availability of data and materials

RNAseq data is available from: the European Nucleotide Archive (ENA) under accession numbers PRJEB56375 (T-RHEX-RNAseq proof-of-principle experiments on MM1.S and HSPCs) and PRJEB47791 (HSPCs) [24]; and the Gene Expression Omnibus (GEO) under accession number GSE173673 (antigen specific CD4 T cells) [23].

### Authors’ contributions

C.G., J.H. and R.M. devised the T-RHEX-RNAseq protocol; J.H., N.F. and S.L. prepared and FACS sorted cells; C.G., N.F. and A.K. performed RNAseq experiments; C.G., J.H., S.L. and R.M. analyzed the data; S.L. and R.M. supervised the research; and all authors contributed to the writing of the manuscript.

## Acknowledgements

We thank: Diagenode for providing un-loaded Tn5 for testing; Prof. Joakim Dillner and colleagues for access to the NextSeq 500 system and the Swedish National Infrastructure for Computing (SNIC) at Uppmax for computational resources.

## SUPPLEMENTAL INFORMATION

Figure S1

Figure S2

Figure S3

Table S1

Working protocol

**Figure S1.**
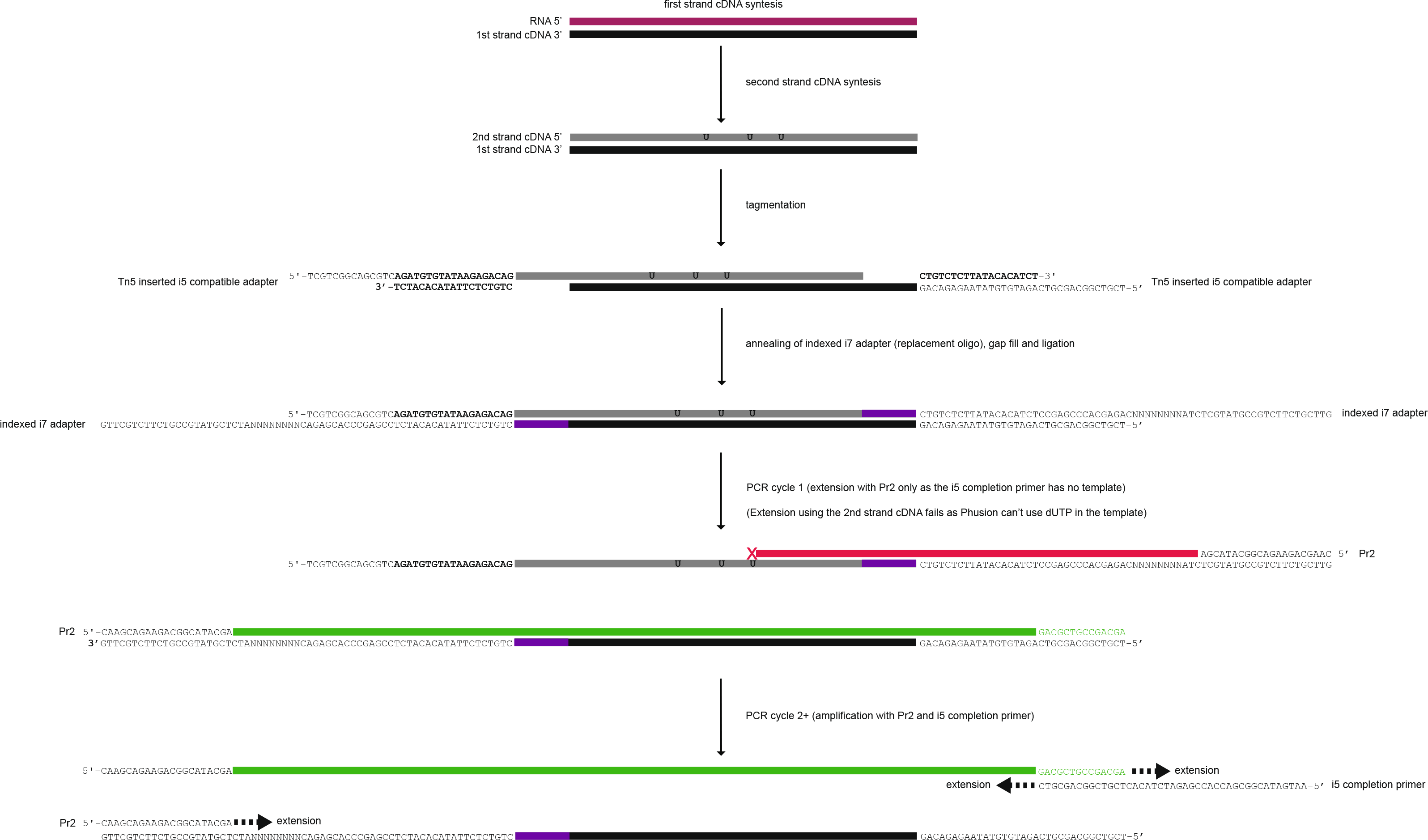
Schematic overview of the T-RHEX-RNAseq protocol outlining adapters, adapter introduction and primers used to amplify the library. In brief, double stranded cDNA reverse transcription with dUTP incorporated during the second strand synthesis is subjected to tagmentation with Tn5 loaded with i5 adapters. The i7 adapters are introduced by annealing an i7 oligo to the covalently attached part of the i5 adapter. Subsequently, gap fill in combination with ligation is used to covalently attach the i7 adapter. As Phusion is unable to utilize the dUTP containing strand as a template, stranded libraries are then generated by amplification using Pr2 in combination with the i5 completion primer.

**Figure S2.**
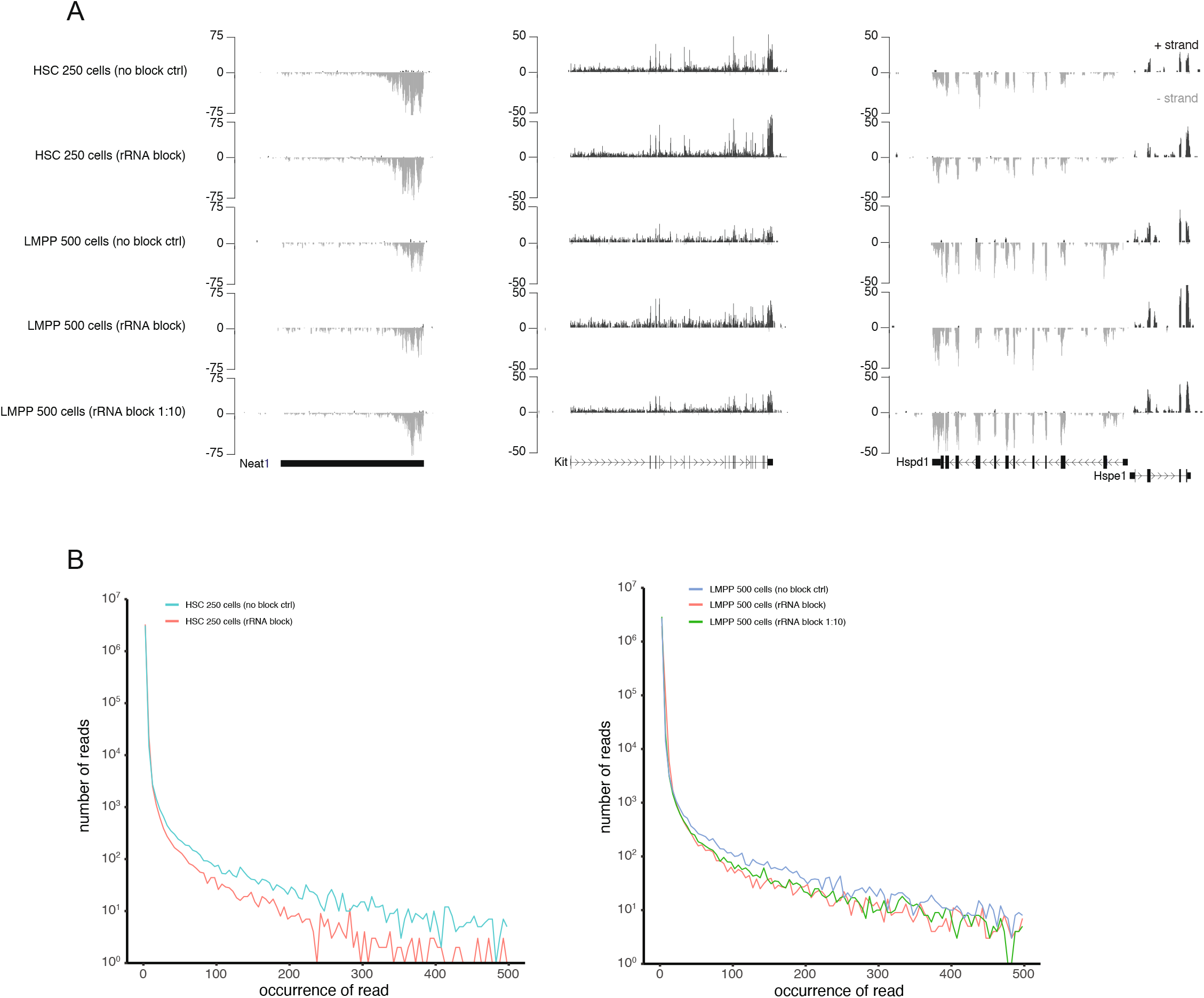
Tracks and duplication rates of T-RHEX-RNAseq libraries from primary hematopoietic stem- and progenitor cells. (A) Tracks showing plus and minus strand reads in the *Neat1*, *Kit*, *Hspd1* and *Hspe1* genomic regions in primary mouse hematopoietic stem cells (HSCs) and lymphoid primed multipotent progenitors (LMPPs). RNAseq libraries were prepared directly from the indicated numbers of cells lysed in Single cell lysis solution (SCLS). The use of rRNA blocking reagents and dilution of the blocking reagent is indicated in parenthesis. (B) Reoccurrence (duplication rates) of reads in the indicated libraries.

**Figure S3.**
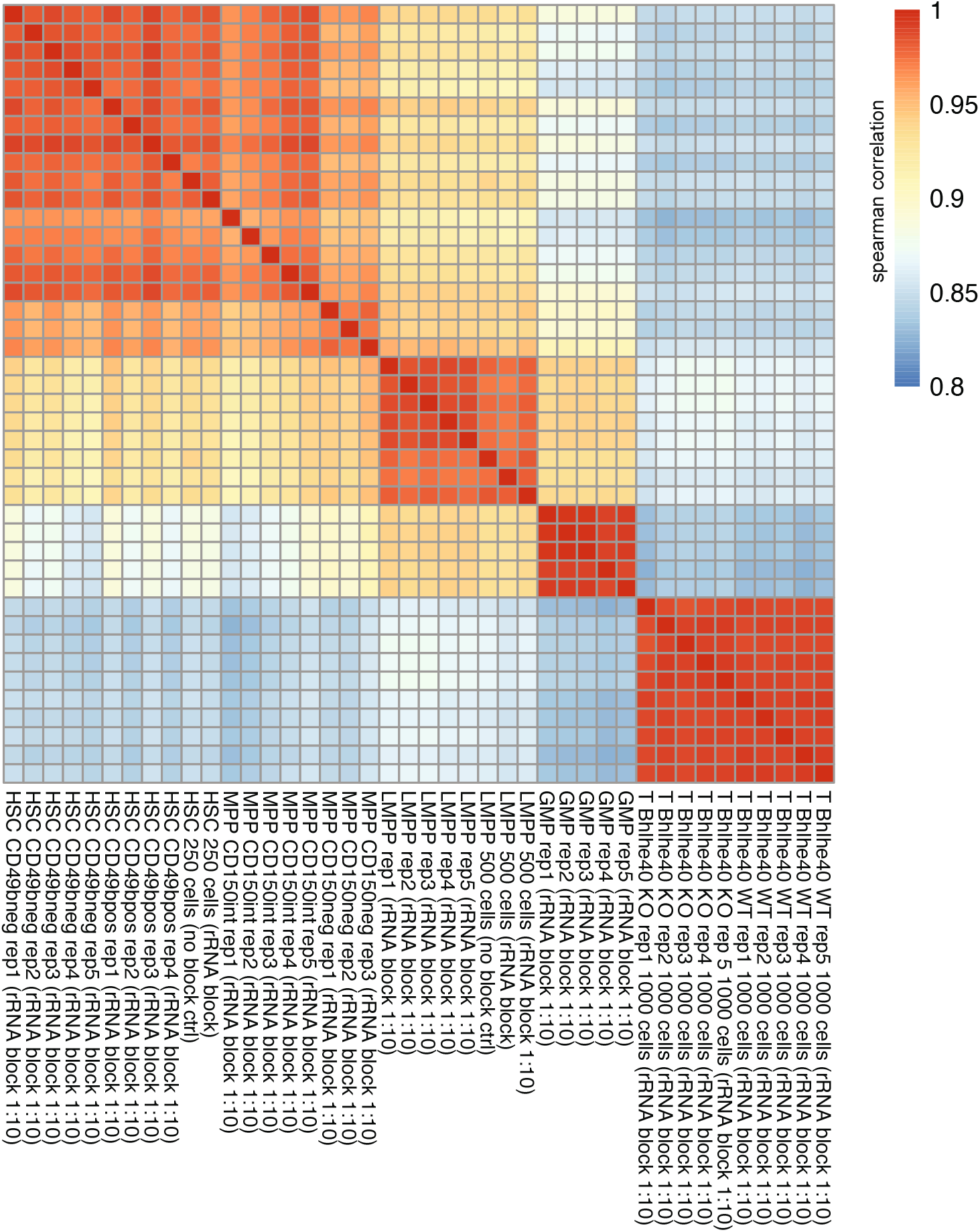
T-RHEX-RNAseq provides highly reproducible data. Spearman correlation between rlog of gene expression in samples generated from the indicated population. The use of rRNA blocking reagents and dilution of the blocking reagent is indicated in parenthesis. Hematopoietic stem cell (HSC with or without CD49b expression); Multipotent progenitor (MPP with no or low CD150 expression), lymphoid primed multipotent progenitors (LMPPs); granulocyte/monocyte progenitors (GMP); and antigen specific CD4 T cells (T, from wild-type or Bhlhe40 knockout mice). Data is from proof-of-principle experiments (HSC and LMPP; 250 and 500 cells respectively) or the subsequently generated T-RHEX-RNAseq data from antigen specific CD4 T cells (1000 cells) [1] and hematopoietic stem/progenitor cells (HSPCs; 250-500 cells) [2]. 1. Rauschmeier R, Reinhardt A, Gustafsson C, Glaros V, Artemov AV, Dunst J, et al. Bhlhe40 function in activated B and TFH cells restrains the GC reaction and prevents lymphomagenesis. J Exp Med. 2021;219:e20211406. 2. Somuncular E, Hauenstein J, Khalkar P, Johansson A-S, Dumral Ö, Frengen NS, et al. CD49b identifies functionally and epigenetically distinct subsets of lineage-biased hematopoietic stem cells. Stem Cell Rep. 2022;17:1546–60.

**Table S1.**
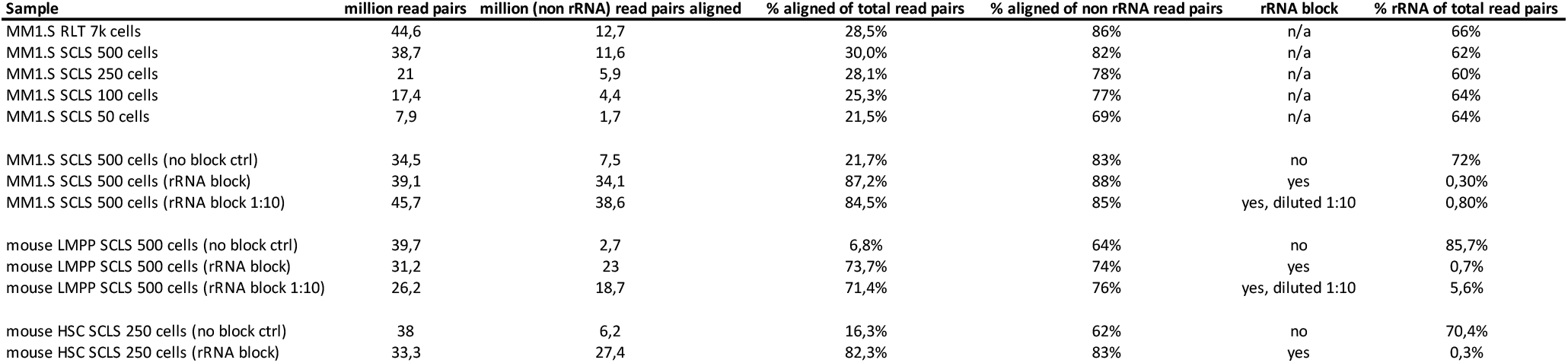
Sample metrics and QC.

| Sample | million read pairs | million (non rRNA) read pairs aligned | % aligned of total read pairs | % aligned of non rRNA read pairs | rRNA block | % rRNA of total read pairs |
| --- | --- | --- | --- | --- | --- | --- |
| MM1.S RLT 7k cells | 44,6 | 12,7 | 28,5% | 86% | n/a | 66% |
| MM1.S SCLS 500 cells | 38,7 | 11,6 | 30,0% | 82% | n/a | 62% |
| MM1.S SCLS 250 cells | 21 | 5,9 | 28,1% | 78% | n/a | 60% |
| MM1.S SCLS 100 cells | 17,4 | 4,4 | 25,3% | 77% | n/a | 64% |
| MM1.S SCLS 50 cells | 7,9 | 1,7 | 21,5% | 69% | n/a | 64% |
| MM1.S SCLS 500 cells (no block ctrl) | 34,5 | 7,5 | 21,7% | 83% | no | 72% |
| MM1.S SCLS 500 cells (rRNA block) | 39,1 | 34,1 | 87,2% | 88% | yes | 0,30% |
| MM1.S SCLS 500 cells (rRNA block 1:10) | 45,7 | 38,6 | 84,5% | 85% | yes, diluted 1:10 | 0,80% |
| mouse LMPP SCLS 500 cells (no block ctrl) | 39,7 | 2,7 | 6,8% | 64% | no | 85,7% |
| mouse LMPP SCLS 500 cells (rRNA block) | 31,2 | 23 | 73,7% | 74% | yes | 0,7% |
| mouse LMPP SCLS 500 cells (rRNA block 1:10) | 26,2 | 18,7 | 71,4% | 76% | yes, diluted 1:10 | 5,6% |
| mouse HSC SCLS 250 cells (no block ctrl) | 38 | 6,2 | 16,3% | 62% | no | 70,4% |
| mouse HSC SCLS 250 cells (rRNA block) | 33,3 | 27,4 | 82,3% | 83% | yes | 0,3% |

## T-RHEX-RNAseq on 250-1000 cells sorted into Single cell lysis solution

**Cell sorting into SCLS** (Single cell lysis kit Invitrogen, Thermo Fisher Scientific cat# 4458235)

1. Use tubes that fit your sorter, for example Biorad 0.2ml flat cap strip tubes (cat# TLS0801 and TCS0803^1^) and make sure the stream hits the lysis solution at the bottom of the tube.
2. Sort up to 1000 cells into tubes containing 0.5μl DNase I and 4.5μl Single cell lysis solution. Keep tubes at 4°C during sorting.
3. Incubate at RT 15min (no mixing is required).^2^
4. Add 0.5μl Single cell stop solution. No mixing is required.
5. Incubate at RT 2min.
6. Snap freeze on dry ice.
7. Transfer to −80°C until further use.

**Primer annealing and rRNA block** (NEBNext Ultra™ II RNA First Strand Synthesis Module, NEB cat#E7771; QIAseq FastSelect -rRNA HMR kit, Qiagen cat# 334386)

Work on ice. In a PCR tube, combine:

5μl Lysed cells
4μl NEBNext 1^st^ strand synthesis reaction buffer (lilac)
1μl NEBNext Random hexamer primers (lilac)
<u>1μl Qiagen FastSelect rRNA HMR reagent^3^</u>
Total 11μl

| PRIME + rRNA-BLOCK |  |
| --- | --- |
| Lid 105°C |  |
| 75°C | 2 min |
| 70°C | 2 min |
| 65°C | 2 min |
| 60°C | 2 min |
| 55°C | 2 min |
| 37°C | 2 min |
| 25°C | 2 min |
| 4°C | hold |

Flick the tube/strip, quickly spin down. Incubate sample in a PCR machine using the PRIME + rRNA-BLOCK program.

**First strand cDNA synthesis** (NEBNext Ultra™ II RNA First Strand Synthesis Module, NEB cat#E7771)

Work on ice. Continuing in the same PCR tube, combine:

11μl primed and blocked RNA (from previous step)
8μl NEBNext Strand specificity reagent (white/brown tube)
<u>2μl NEBNext First strand synthesis enzyme mix (lilac)</u>
Total 21μl

| cDNA1STR |  |
| --- | --- |
| Lid 105°C |  |
| 25°C | 10 min |
| 42°C | 50 min |
| 70°C | 15 min |
| 4°C | hold |

Flick the tube/strip, quickly spin down. Incubate sample in a PCR machine using the cDNA1STR program.

**Second strand cDNA synthesis** (NEBNext Ultra™ II Directional RNA Second Strand Synthesis Module, NEB Cat# E7550)

Work on ice. Continuing in the same PCR tube combine:

21μl cDNA sample (from previous step)
8μl NEBNext 2^nd^ strand synthesis reaction buffer dUTP (orange)
4μl NEBNext 2^nd^ strand synthesis enzyme mix (orange)
<u>48μl nuclease free water</u>
Total 81μl

| cDNA2STR |  |
| --- | --- |
| Lid 40°C |  |
| 16°C | 60 min |
| 80°C | 15 min |
| 4°C | hold |

Flick the tube/strip, quickly spin down. Incubate sample in a PCR machine using the cDNA2STR program.

(Safe stopping point. Samples can be kept overnight at 4°C.)

**Tagmentation** (using Tn5 enzyme containing only i5 adapters)

Work on ice. Continuing in the same PCR tube combine:

81μl double stranded (ds) cDNA sample (from previous step)
20μl 5X Tn5 Tagmentation buffer (home-made, see reagents section)
<u>1μl Tn5 with i5 adapters (see reagents section)</u>
Total 102μl

|  |  |
| --- | --- |
| dscDNATN5 |  |
| Lid 105°C |  |
| 55°C | 5 min |
| 4°C | hold |

Flick the tube/strip, quickly spin down. Incubate sample in a PCR machine using the dscDNATN5 program.

Add 5μl Tagmentation Stop Solution (1% SDS, see reagents section), mix and quickly spin down. Incubate at RT 5min.

**Bead cleanup of library** (Ampure XP beads, Beckman Coulter cat#A63881; DynaMag™-96 Side Magnet, Thermo Fisher Scientific cat#12331D)

Let beads adjust to RT for 15-30min.

1. Mix Ampure beads thoroughly, add 130μl beads (1.2:1) to sample (107μl). Set pipette to 230μl and pipette x10 in tube. Incubate at RT for 5 min.
2. Place sample on magnet for 5 min until supernatant is clear.
3. Gently remove and discard 235μl of the supernatant without disturbing the beads.
4. With sample still on magnet, add 200μl 80% EtOH, leave for 30s, then gently discard all supernatant. Repeat once for a total of two washings.
5. Quickly spin down tube, put back on magnet and remove all remaining EtOH. (Optionally, if not using a spindown, let beads air dry for 3 min.) Do not let beads over-dry.
6. Resuspend sample in 15μl water, incubate at RT for 2 min.
7. Place sample on magnet for 3-5 min until supernatant is clear.
8. Transfer 14μl of clear supernatant to a new PCR tube (preferably containing index and gap fill reagents, see next step below).

### Oligo replacement and gap fill

Thaw oligos and buffer at RT and keep on ice. Gap fill buffer and Replacement index can be combined in a new tube during the above Ampure cleanup. Then add supernatant directly into tube with reagents.

14μl tagmented sample (from previous step)
4μl Gap fill buffer (home-made, see reagents section)
<u>1μl Replacement index i7 oligo (40μM, index Ad2.1-Ad2.24, see reagents section)</u>
Total 19μl

Pipette or flick the tube/strip, quickly spin down. Incubate sample in a PCR machine using the RNAGF program.

|  |  |
| --- | --- |
| RNAGF |  |
| Lid 105°C |  |
| 45°C | 1 min |
| 37°C | 30 min |
| PAUSE |  |
| 37°C | 30 min |
| 4°C | hold |

After 30 min pause program and add:

<u>1.5μl Gap fill and ligation enzyme mix (see reagents section)</u>
Total 20.5μl

No mixing is required. Continue incubation for 30 more minutes.

**Bead cleanup** (Ampure XP beads, Beckman Coulter cat#A63881; DynaMag™-96 Side Magnet, Thermo Fisher Scientific cat#12331D)

Let beads adjust to RT for 15-30min.

1. Mix Ampure beads thoroughly, add 25μl beads (1.2:1) to sample (20.5ul). Set pipette to 40μl and pipette x10 in tube. Incubate at RT for 5 min.
2. Place sample on magnet for 5 min until supernatant is clear.
3. Gently remove and discard 44μl of the supernatant without disturbing the beads.
4. With sample still on magnet, add 200μl 80% EtOH, leave for 30s, then gently discard all supernatant. Repeat once for a total of two washings.
5. Quickly spin down tube, put back on magnet and remove all remaining EtOH. (Optionally, if not using a spindown, let beads air dry for 3 min.) Do not let beads over-dry.
6. Resuspend sample in 13μl water, incubate at RT for 2 min.
7. Place sample on magnet for 3-5 min until supernatant is clear.
8. Transfer 12μl of clear supernatant to a new PCR tube (preferably containing library PCR amplification reagents, see next step below).

### Library PCR amplification

Thaw reagents at RT and then keep on ice. Reagents can be combined in a new tube during the Ampure cleanup. Then add supernatant directly into tube with reagents.

12μl sample (from previous step)
0.5μl PR2+Ad1 primer (10μM, see reagents section)
<u>12.5μl 2X Phusion HF PCR Master mix (NEB cat#M0531)</u>
Total 25μl

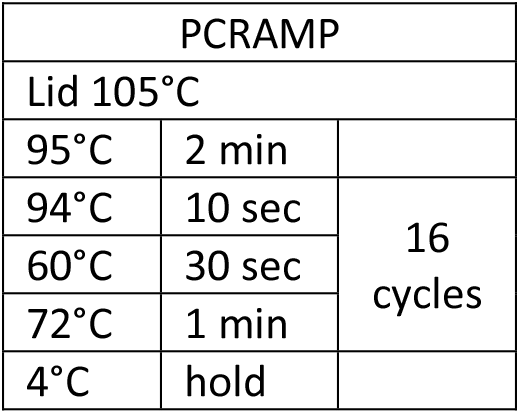

Pipette or flick the tube/strip, quickly spin down. Incubate sample in a PCR machine using the PCRAMP program.

**Post amplification bead cleanup** (Ampure XP beads, Beckman Coulter cat#A63881, DynaMag™-96 Side Magnet Thermo Fisher Scientific cat#12331D)

Let beads adjust to RT for 15-30min.

1. Mix Ampure beads thoroughly, add 23μl beads (0.9:1) to sample (25ul), Set pipette to 40μl and pipette x10 in tube. Incubate at RT for 5 min.
2. Place sample on magnet for 5 min until supernatant is clear.
3. Gently remove and discard 46μl of the supernatant without disturbing the beads.
4. With sample still on magnet, add 200μl 80% EtOH, leave for 30s, then gently discard all supernatant. Repeat once for a total of two washings.
5. Quickly spin down tube, put back on magnet and remove all remaining EtOH. (Optionally, if not using a spindown, let beads air dry for 3 min.) Do not let beads over-dry.
6. Resuspend sample in 13μl water, incubate at RT for 2 min.
7. Place sample on magnet for 3-5 min until supernatant is clear.
8. Transfer 11-12μl of clear supernatant to a new PCR tube.

Quantify and quality check libraries with Qubit (Qubit™ dsDNA HS Assay Kit, Thermo Fisher Scientific cat# Q32851) and TapeStation (TapeStation DNA Screen tape HSD1000, Agilent cat#5067-5584).

Store samples at −20°C until sequencing.^4^

### Reagents; kits/commercial

Single cell lysis kit Invitrogen (Thermo Fisher Scientific cat# 4458235)
NEBNext Ultra™ II RNA First Strand Synthesis Module (New England Biolabs cat#E7771)
NEBNext Ultra™ II Directional RNA Second Strand Synthesis Module (New England Biolabs cat# E7550)
QIAseq FastSelect -rRNA HMR kit (Qiagen cat# 334386)
Agencourt Ampure XP beads (Beckman Coulter cat#A63881)
2X Phusion High-Fidelity PCR Master Mix with HF Buffer (New England Biolabs cat#M0531)
DynaMag™-96 Side Magnet (Thermo Fisher Scientific cat#12331D)
Qubit™ dsDNA HS Assay Kit (Thermo Fisher Scientific cat# Q32851)
TapeStation DNA Screen tape HSD1000 (Agilent cat#5067-5584)

### Reagents; home-made or home-mixed

#### 5X Tn5 Tagmentation buffer

Total volume=2ml for 200 samples, aliquot in 4×500μl.

|  |  |
| --- | --- |
| 50mM Tris acetate pH7.6 | 100µl (from 1M stock) |
| 25mM Mg acetate | 50µl (from 1M stock) |
| 50% dimethylformamide | 1000µl (from 100% stock, add DMFA in a hood) |
| H2O | 850µl |

#### Tn5 enzyme containing only i5 adapters

We have either used home-made or commercially available Tn5, loaded with only the i5 adapter. The home-made Tn5 (kindly provided by Prof. Richard Sandberg) was diluted to 15μM in Tn5 storage buffer before use.

Alternatively, unloaded tagmentase (Tn5 transposase, Diagenode cat#C01070010) was loaded with i5 adapters. Adapter annealing and loading was performed following the manufacturer’s protocol (https://www.diagenode.com/files/protocols/PRO-Transposome-Assembly-V2.pdf) but using Tn5MErev annealed to i5MEadapt (see below) and adding twice the amount of this annealed product into the Tn5 adapter loading reaction. Before use, the Tn5 was diluted 1:4, 1:8 or 1:16 in Tn5 storage buffer with similar results.^5^

Tn5 i5 adapters

Tn5MErev: 5’-[phos]**CTGTCTCTTATACACATCT**-3’ annealed to:
i5MEadapt: 5’-TCGTCGGCAGCGTC**AGATGTGTATAAGAGACAG**-3’

#### Tn5 storage buffer

|  |  |
| --- | --- |
| 50 mM HEPES, pH 7.2 | 5 ml 1M |
| 100 mM NaCl | 2 ml 5 M |
| 0.1 mM EDTA | 20 µl 500 mM |
| 1 mM DTT | 0.1 ml 1M |
| 0.1% triton X-100 | 1 ml 10% |
| 50% glycerol | 50ml 100% |
| H2O | 42.8ml |

**Tagmentation Stop Solution**, 1% SDS in water.

#### Gap fill buffer

Total volume=1ml for 250 samples, filter, and aliquot in 4×250μl.

|  |  |
| --- | --- |
| 165mM tris acetate pH7.5 | 165µl (from 1M stock RT) |
| 330mM K acetate | 330µl (from 1M stock RT) |
| 50mM Mg acetate | 50µl (from 1M stock RT) |
| 2.5mM DTT | 25µl (from 100mM stock -20°) |
| 1.25mM dATP | 12.5µl (from 100mM stock -20°) |
| 1.25mM dCTP | 12.5µl (from 100mM stock -20°) |
| 1.25mM dGTP | 12.5µl (from 100mM stock -20°) |
| 1.25mM dTTP | 12.5µl (from 100mM stock -20°) |
| 1mM beta-NAD | 26.5µl (from 37.7mM stock: 25mg+ 1ml H<sub>2</sub>O -20°) |
| H<sub>2</sub>O | 353.4µl |

#### Replacement index i7 oligo (Ad2.index)

Replacement oligos (custom made by IDT, ordered in 100nmole scale) diluted to 40μM working concentration in water. (* indicates phosphothioate bonds between the last 8 bases.)

Ad2.index 5’-[phos]CTGTCTCTTATACACATCTCCGAGCCCACGAGAC<u>NNNNNNNN</u>ATCTCGTATGCCGTCT T*C*T*G*C*T*T*G

|  |  |  |  |
| --- | --- | --- | --- |
| Ad2.1 | TAAGGCGA | Ad2.13 | GTCGTGAT |
| Ad2.2 | CGTACTAG | Ad2.14 | ACCACTGT |
| Ad2.3 | AGGCAGAA | Ad2.15 | TGGATCTG |
| Ad2.4 | TCCTGAGC | Ad2.16 | CCGTTTGT |
| Ad2.5 | GGACTCCT | Ad2.17 | TGCTGGGT |
| Ad2.6 | TAGGCATG | Ad2.18 | GAGGGGTT |
| Ad2.7 | CTCTCTAC | Ad2.19 | AGGTTGGG |
| Ad2.8 | CAGAGAGG | Ad2.20 | GTGTGGTG |
| Ad2.9 | GCTACGCT | Ad2.21 | TGGGTTTC |
| Ad2.10 | CGAGGCTG | Ad2.22 | TGGTCACA |
| Ad2.11 | AAGAGGCA | Ad2.23 | TTGACCCT |
| Ad2.12 | GTAGAGGA | Ad2.24 | CCACTCCT |

#### Gap fill and ligation enzyme mix

Sulfolobus DNA Polymerase IV 100 units (2U/μl) 50μl NEB (Bionordika cat# M0327S) E. coli DNA Ligase 1000U (10U/μl) 100μl NEB (Bionordika cat# M0205L)

Mix 10μl of polymerase with 20μl of ligase (Unit ratio 1:10), store at −20°C.

#### PR2+Ad1 primer mix

10μl of Primer 2 (PR2, 100μM stock in water), 10μl Adapter 1 (Ad1, 100μl stock in water) and 80μl water was mixed to make the 10μM PR2+Ad1 primer mix working solution.

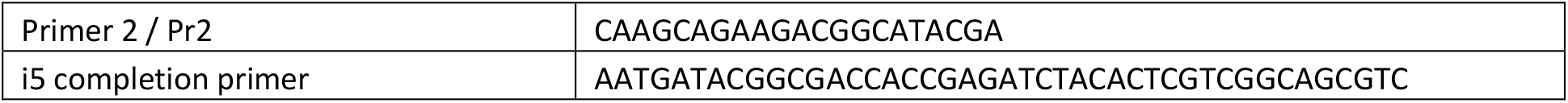

## Footnotes

1. Tubes can be individually fastened to 96-well racks (Biorad cat#TRC9601) for easy handling in plate format.

2. We use a prolonged incubation at RT compared to kit protocol, to reduce DNA content seen in some samples sorted with cooling.

3. 1:10 dilution of HMR reagent in water gives similar results.

4. Paired-end 41sequencing using a Nextseq500/550 75 cycle or Nextseq2000 P3 50 cycle kit is an affordable option compatible with the insert size of the library.

5. Likely the Tn5 can be further diluted without impacting the results.

